# The downward spiral: eco-evolutionary feedback loops lead to the emergence of ‘elastic’ ranges

**DOI:** 10.1101/008458

**Authors:** Alexander Kubisch, Anna-Marie Winter, Emanuel A. Fronhofer

**Affiliations:** Institute for Landscape and Plant Ecology, University of Hohenheim, August-von-Hartmann-Str. 3, 70599 Stuttgart, Germany; Institute des Sciences de l’Evolution, UMR 5554 | CNRS, Université Montpellier II, Place Eugène Bataillon, 34095 Montpellier, France; Department of Biosciences, Centre for Ecological and Evolutionary Synthesis (CEES), Postboks 1066 Blindern, 0316 Oslo, Norway; Eawag: Swiss Federal Institute of Aquatic Science and Technology, Department of Aquatic Ecology, Überlandstrasse 133, CH-8600 Dübendorf, Switzerland

**Keywords:** elastic ranges, Allee effects, eco-evolutionary dynamics, competition, dispersal evolution, range dynamics, individual-based model

## Abstract

In times of severe environmental changes and resulting shifts in the geographical distribution of animal and plant species it is crucial to unravel the mechanisms responsible for the dynamics of species’ ranges. Without such a mechanistic understanding, reliable projections of future species distributions are difficult to derive. Species’ ranges may be highly dynamic. One particularly interesting phenomenon is range contraction following a period of expansion, referred to as ‘elastic’ behaviour. It has been proposed that this phenomenon occurs in habitat gradients, which are characterized by a negative cline in selection for dispersal from the range core towards the margin, as one may find, for example, with increasing patch isolation. Using individual-based simulations and numerical analyses we show that Allee effects are an important determinant of range border elasticity. If only intra-specific processes are considered, Allee effects are even a necessary condition for ranges to exhibit elastic behavior. The eco-evolutionary interplay between dispersal evolution, Allee effects and habitat isolation leads to lower colonization probability and higher local extinction risk after range expansions, which result in an increasing amount of marginal sink patches and consequently, range contraction. We also demonstrate that the nature of the gradient is crucial for range elasticity. Gradients which do not select for lower dispersal at the margin than in the core (especially gradients in patch size, demographic stochasticity and extinction rate) do not lead to elastic range behavior. Thus, we predict that range contractions are likely to occur after periods of expansion for species living in gradients of increasing patch isolation, which suffer from Allee effects.

## Introduction

Currently, a significant number of species’ ranges have come under pressure through global climatic, environmental and socio-economic changes (Walther et al., 2002, Perry et al., 2005, Chen et al., 2011, Bellard et al., 2013). It is now well documented that such range shifts often exhibit subsequent fundamental shifts in (meta-)population dynamics (Altermatt et al., 2008, Thomas, 2010). If we intend to assess and predict the impact of these global changes (Stocker et al., 2013) an adequate understanding of range dynamics is crucial. Yet, our understanding of the formation of species’ ranges is still limited, as both biotic and abiotic ecological, as well as rapid evolutionary, processes affect range formation in complex ways (Kubisch et al., 2014, Fronhofer and Altermatt, 2015).

Particularly important and complex are non-equilibrium situations, in which species’ ranges either expand or contract. In this respect, the dynamics of range expansions have been extensively studied theoretically, but also the comparative and experimental literature is growing. From a purely ecological perspective, range expansions have been described by reaction-diffusion models (Fisher, 1937, Kolmogorov et al., 1937, Lubina and Levin, 1988). Melbourne and Hastings (2009) demonstrated experimentally the intrinsic role of stochasticity during range expansions. These results are underpinned by the work of Giometto et al. (2014), who showed that this stochasticity can indeed be predicted. However, range expansions rarely only include ecological dynamics. It is now clear that rapid evolutionary and resulting eco-evolutionary dynamics may play a major role (Kubisch et al., 2014, Perkins et al., 2013, Fronhofer and Altermatt, 2015). For example, high emigration rates are selected for during the process of expansion due to spatial (Phillips et al., 2010, Shine et al., 2011, Fronhofer and Altermatt, 2015, more dispersive individuals being at the front in combination with fitness benefits through reduced competition;) and kin selection (Kubisch et al., 2013) which leads to accelerating expansions. From a genetic point of view, range expansions usually lead to decreasing genetic diversity, either affecting the adaptation of species (Excoffier et al., 2009) or their dispersal ability directly (Cobben et al., 2015). In addition, expanding populations are also likely to suffer from a mutation load, also called expansion load (Peischl et al., 2015). Finally, Allee effects, which can be the consequence of e.g. sexual reproduction or sociality Courchamp et al. (2010), can drastically influence expansion dynamics, as they can lead to pulsed patterns of invasions (Johnson et al., 2006, Schurr et al., 2008).

Range contractions, however, are more difficult to investigate empirically and thus have been less intensively studied (Channell and Lomolino., 2000). Yet, range contractions are highly relevant from a conservation and management point of view as they are usually assumed to be caused by extrinsic mechanisms, like climate change or human impacts (Li et al., 2015).

To further complicate the situation, besides simply expanding or contracting, species’ ranges may also exhibit more complex dynamics such as ‘elastic’ behavior. Elasticity (as described by Kubisch et al., 2010) implies that a range expansion is immediately followed by a period of contraction due to evolutionary changes in dispersal. In his review of the work of MacArthur (1972), Holt (2003) first described this phenomenon. He argued that after a period of increasing dispersal during range expansion there can be substantial selection against dispersal in marginal areas due to source-sink dynamics. If invasions occur along a gradient from source to sink populations, the latter would be sustained by initially high emigration rates which are typical for such expansions (Shine et al., 2011). Subsequent selection against dispersal due to an increased probability of arriving in sink patches characterized by low fitness expectations will result in a contraction of the geographical range.

In a simulation study, Kubisch et al. (2010) could show that this phenomenon may indeed be likely to occur in nature, but that it crucially depends on underlying gradient. The authors found that range border elasticity could only be observed in fragmentation gradients and, to a smaller extent, in fertility gradients. They concluded that the mechanism explaining range elasticity is selection for lower emigration rates at range margins relative to core areas. In more recent work Henry et al. (2013) suggested that under climate change, elasticity should also be found in gradients of patch size, habitat availability, growth rate and local extinction risk.

Following the argumentation by Holt (2003) and MacArthur (1972), a crucial determinant of range border elasticity is the presence of actual sink patches at the initial wide range after expansion. Sink populations are populations with a negative growth rate and may be the result of altered abiotic conditions, which lead to maladaptation and reduced growth. Similarly, sink patches may be caused by new or altered biotic interactions, such as the occurrence of a predator at the range margin. Yet, previous studies (Kubisch et al., 2010, Henry et al., 2013) report the occurrence of range border elasticity even without changes in abiotic local conditions or the occurrence of novel biotic interactions. Therefore, intra-specific, biotic processes must be sufficient to generate sink patches at the range margin. Under these conditions an important mechanism leading to the emergence of sinks are demographic Allee effects, which are defined as reduced growth rates at low population sizes or densities in comparison to populations at intermediate densities (Courchamp et al., 2010).

Here we argue that a negative cline in selection for dispersal from the range core to the margin is only one prerequisite for range elasticity caused by intraspecific processes, and that the presence of Allee effects leading to sink populations at range margins is the second. The eco-evolutionary feedback loop created by these two forces leads to a spatio-temporally non-linear cline in immigrant fitness which is caused by the emergence of sink patches and finally results in range contraction.

## The Model

We use an individual-based model of a spatially structured population of an asexually reproducing species with discrete generations. This approach has been used in several previous studies (Poethke et al., 2011, Fronhofer et al., 2013, Kubisch et al., 2014).

### Landscape

We implement linear unidirectional environmental gradients. This means that along the *x*-axis of the landscape, one specific habitat characteristic changes from favorable to unfavorable conditions with respect to the survival of the species (see below for details). The simulated landscape consists of *x* ⋅ *y* = 200 ⋅ 50 habitat patches, arranged on a rectangular grid. Larger landscapes (specifically in *x*-direction) would lead to larger elasticity effects. Yet, we were computationally limited in the total number of populations.

### Individuals

Every patch may contain a population of the species, assuming a carrying capacity *K*_*x,y*_(see below). Local populations consist of individuals, which are determined by their specific location *x, y* and one heritable trait defining their probability to emigrate.

### Population dynamics

Local population dynamics follow the discrete logistic growth model developed by Beverton and Holt (1957). This model is extended by the implementation of a direct Allee effect, the strength of which depends on population density instead of size (see also Kubisch et al., 2011). We draw the individuals’ average offspring number for every patch and generation 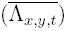 from a log-normal distribution with mean *λ*_*x,y*_ and standard deviation *σ*. The latter thus represents the degree of environmental stochasticity. Afterwards every individual in a patch gives birth to a number of offspring drawn from a Poisson distribution with mean 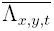. Density-dependent competition then acts on offspring survival probability *s*, which is given by

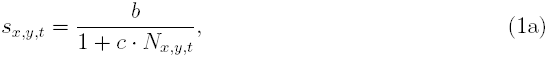

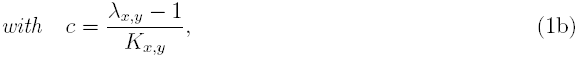

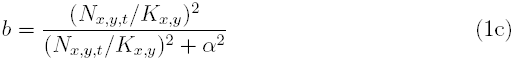

with *K*_*x,y*_ being carrying capacity of a patch, *N*_*x,y,t*_ denoting the population size of a focal patch and *α* defining the strength of the Allee effect. Note that propagule survival is thus actually regulated by the number of parental individuals, the competition between which is thus phenomenologically implemented. We assume a sigmoid increase in survival probability with the number of inhabitants in a patch (see eq. 1c). Generally, increasing *α* leads to a decreased probability of survival. For example, individuals in a population of density 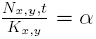 will have a decrease in their survival of 50 %.

A newborn inherits the dispersal allele from its parent. During this process the allele may mutate with probability *m* = 10^−4^. In case of a mutation, a Gaussian distributed random number with mean 0 and standard deviation 0.2 is added to the allele’s value. As has been shown by Kubisch et al. (2010), range limit elasticity is robust against changes in mutation rate, as long as increasing dispersal during range expansion is provided.

We assume a moderately low level of population turnover as is characteristic for metapopulations (Fronhofer et al., 2012). Here, this turnover is driven extrinsically due to local patch extinctions. Therefore, following reproduction, every population may go extinct by chance with probability *ϵ* = 0.05. Changing this extinction rate did in previous analyses not result in qualitative changes of the presented results.

### Dispersal

Surviving offspring may emigrate from their natal patch. We implemented dispersal in two different modes.

#### Nearest-neighbor dispersal

In this standard scenario, which is used throughout the main text, the probability to disperse for any given offspring is given by its dispersal allele (*e*). If an individual disperses, it may die with a certain probability *µ*, which includes all potential costs that may be associated with dispersal, like predation risk or energetic costs (Bonte et al., 2012). The dispersal mortality is calculated as the arithmetic mean between the patch-specific dispersal mortalities *µ*_*x,y*_ of the natal and the target patch. The target patch is randomly drawn from the eight surrounding patches. To avoid edge effects we wrap the world in *y*-direction, thus forming a tube along the *x*-dimension of the world. Individuals leaving the world along the *x*-dimension are reflected.

#### Dispersal kernel

To test the validity of our results against alternative implementations of dispersal we performed additional simulations with dispersal distances evolving instead of propensities. In these cases the dispersal alleles coded for the mean of a negative exponential distance distribution (kernel; see also Henry et al., 2013). Given that the dispersal mortality *µ* in our original approach means a per-step mortality (as the step length for nearest neighbor dispersal is one) we have based the implementation of mortality in the kernel scenario on the same rationale and assumed that the probability of dying during the transition phase, *µ*_*d*_, is given by:

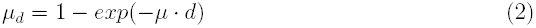

with *d* denoting the traveled dispersal distance (for more details see Fronhofer et al., 2015). *µ* is calculated as described above as the mean of *µ*_*x,y*_ of the natal and the target patch.

### Simulation experiments and analysis

In order not to bias the results by using artificial initial values for the dispersal traits we implemented a burn-in period allowing for the adaptation of dispersal strategies to local conditions in the range core. Therefore we added additional 10 rows to the landscape in front of the position *x* = 1, all patches there being defined by conditions found at position *x* = 1 and filled these patches with *K*_1,*y*_ individuals. We then let the simulation run for 2, 500 generations, assuming torus conditions of the burn-in region (i.e. individuals leaving this region in *x*- and *y*-direction were wrapped around). In the case of dispersal propensities evolving, the alleles were initially drawn from a uniform distribution between 0 and 1. In the alternative scenario with dispersal distances evolving we initialized the individuals with mean dispersal distance values from a uniform distribution between 0 and 10. The fact that dispersal distances did not closely approach the maximum values indicates that our burn-in region was sufficiently large.

After the burn-in phase (i.e. in generation 2,501), we allowed the populations to spread further than the first 10 rows for 5,000 generations, allowing for range expansion and assuring the formation of a stable range limit. Although we focus on a gradient in dispersal mortality (i.e. habitat fragmentation), we tested a range of other possible gradients. A summary of gradient dimensions and references to according figures are given in table 1.

**Table 1:** Applied dimensions of gradients including references to the according results figures. Figures A1 to A10 can be found in the Supplemental Material.

| gradient | favorable condition | unfavorable condition | meaning | results figure |
| --- | --- | --- | --- | --- |
| $\mu$ | 0 | 1 | dispersal mortality | Figs. 1, 2, 3, A6 |
| $K$ | 100 | 0 | carrying capacity | Figs. A1, A7 |
| $\lambda$ | 4 | 0 | growth rate | Figs. A2, A8 |
| $\sigma$ | 0 | 10 | environmental stochasticity | Figs. A3, A9 |
| $\epsilon$ | 0 | 1 | extinction rate | Figs. A4, A10 |

Respective parameters, which were not changing across space in a given simulation were set to standard values (*K*_*x,y*_ = 100, *λ*_*x,y*_ = 2, *µ*_*x,y*_ = 0.2, *σ*_*x,y*_ = 0.2, *ϵ*_*x,y*_ = 0.05). To account for the fact that fragmentation gradients, as they occur in nature, usually affect not only the isolation of habitat patches, but also imply decreasing patch sizes, we have additionally tested a ’mixed’ gradient, in which along the x-axis habitat capacity *K* was reduced and dispersal mortality *µ* increased, using the same parameters as given above. These results can be found in the Supplementary material, Appendix 1, Fig. A5. We repeated the simulations for the five standard gradients (i.e. *µ*, *K*, *λ*, *σ*, *ϵ*) for a scenario with dispersal distance instead of propensity evolution. The results are provided in the Supplementary material, Appendix 2, Fig. A6 - A10). We tested 11 values for the strength of the Allee effect (*α*) in equidistant steps from 0 to 0.1. For all scenarios we performed 50 replicate simulations. For all runs we assessed the absolute range border position *R*, defined as the x-position of the foremost populated patch. We analyzed the marginal emigration rate as the fraction of individuals emigrating from their natal patch in the five columns of patches behind the range border.

To quantify the presence and degree of range elasticity, we analyzed range border position as a function of time. We fitted a function to the resulting progression of relative range size *r*, which is calculated as the absolute range size position along the landscape’s *x*-dimension *R* divided by the maximum extent of that dimension (*x*_*max*_ = 200). The function we used (eq. 3) is flexible enough to quantify elasticity and its parameters can be directly interpreted in biological terms:

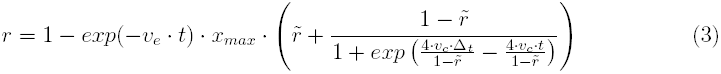

with *v*_*e*_ denoting the speed of range expansion, 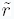 the equilibrium range size, *v*_*c*_ the speed of range contraction and ∆_*t*_ the time to reach equilibrium. Using non-linear least squares regressions (R language for statistical computing version 3.1.0 function nls(), R Core Team, 2013) we fit the curve to the respective simulation output (i.e. relative border position over time). The relative amplitude of the elasticity effect was calculated using the resulting function as:

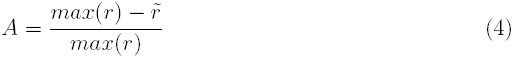

Figure 3E shows a typical example of this calculation.

### Numerical analysis of the mean fitness of colonizers

To mechanistically investigate the eco-evolutionary feedback loop underlying the observed range dynamics, we performed additional numerical analyses. We quantified the mean fitness expectations (reproductive success) of potential colonizers at the range margin as a function of their dispersal strategy and of the landscape gradient. For a better comparability, we calculate the colonizers’ fitness expectations over one generation for every potential location of the range margin in close analogy to the individual-based simulation model described above:

1. We calculate the number of colonizers at the margin based on the conservative assumption that the range behind the margin is fully populated (i.e. all patches are at *N*_*x,y,t*_ = *K*_*x*_). Thus the number of colonizers is given by *N*_*c,x*_ = *K*_*x*_ ⋅ e ⋅ (1 − *µ*_*x*_), with *e* denoting emigration rate. Our approach is conservative in the sense that populations directly behind the margin may not yet have reached carrying capacity. We therefore overestimate the number of colonizers, which leads to a systematic underestimation of the impact of the phenomenon of interest, the Allee effect.
2. As described above for the evolutionary simulations we assume that the mean fecundities of the colonizers 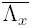 follow a lognormal distribution with mean *λ*_*x*_ and standard deviation *σ*_*x*_. The parameter *σ* accounts for environmental stochasticity.
3. Demographic stochasticity is taken into account by assuming that reproduction can be described as a Poisson process with 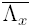 as mean. Subsequently, density regulation is applied according to eq. 1 (with *N*_*c,x*_ as the population size).
4. Finally, local extinctions are represented as a binomial process with mean (1 − *ϵ*) acting on the offspring numbers.

This algorithm allows us to calculate a distribution *W* of potential per capita offspring numbers for each *x*-location in all gradients, assuming that the margin of a saturated range would lie directly at its front. We approximated this distribution by sampling 1, 000, 000 times. To average the mean colonizer fitness we used an approximated geometric mean calculation. The arithmetic mean would be a poor estimate of fitness, as it is comparably insensitive to the distribution’s variance. The true geometric mean is, however, too sensitive against zero offspring numbers, which are drawn with a high probability based on above method. A Jean series approximation of the geometric mean, however, provides a sensible estimate, as it is sensitive towards variation but not zero, when such offspring numbers are included. According to Jean and Henry (1983) we calculate the mean colonizer fitness 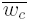 thus as:

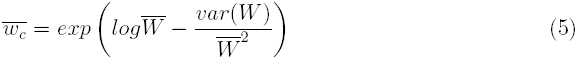

with 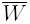 denoting the arithmetic mean of the distribution. In summary, *w*_*c*_ thus is the average number of offspring an immigrant would get, if it would colonize a habitat patch at the very margin of the species’ distribution. This metric depends strongly on the dispersal rate, local environmental conditions and the strength of the Allee effect. We performed the analysis for all gradients and two values of Allee effect strength (*α* = 0 and *α* = 0.05). We varied emigration rate *e* in 10 equidistant steps from 0.05 to 0.5.

## Results

Strong elasticity is only detected for a gradient in dispersal mortality, i.e. patch fragmentation, in the presence of an Allee effect (Fig. 1). We found a very weak elastic behavior in a gradient of decreasing per capita growth rate (*λ*_0_; Supplementary material Appendix 1, Fig. A2) — a result consistent with the findings of Kubisch et al. (2010). All other gradients (*K*, *σ*, *ϵ*) do not lead to range border elasticity (Supplementary material Appendix 1, Fig. A1, A3, A4). Importantly, elasticity only occurs in the presence of an Allee effect and the degree of elasticity (amplitude) increases with increasing Allee effect strength (Fig. 1,3). For values of *α* exceeding 0.07, the entire spatially structured population goes extinct. It is also important to note that we did find elastic ranges for a mixed gradient - scenario, including declining patch size and increasing patch isolation (Supplementary material Appendix 1, Fig. A5).

**Figure 1:**
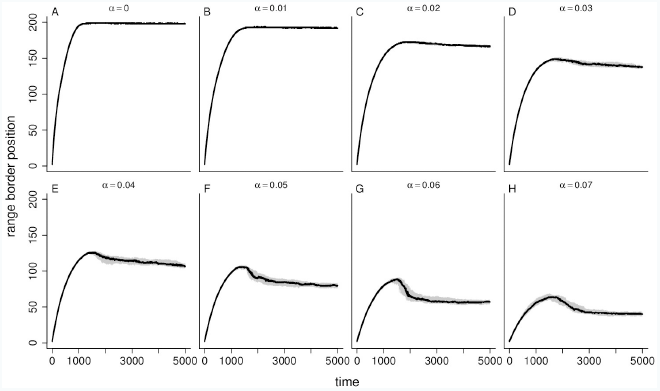
Range border position as a function of simulation time for a gradient in dispersal mortality (*µ*). Allee effect strength increases from A to H. For parameter values see main text. The black lines show the median values of 50 replicate simulations, the shaded grey areas denote 25% – and 75% quantiles.

The evolving emigration rate at the range margin is negatively affected by the Allee effect strength (Fig. 2). Consequently, with increasing Allee effects range expansion is slower (Fig. 3A) and the resulting equilibrium range size is smaller (Fig. 3B). The enhanced elasticity for increasing Allee effects (which is apparent in Fig. 1) is characterized by (i) an increase in the velocity of contraction (followed by a slight decrease for a very strong Allee effect; Fig. 3C) and (ii) an increase in the amplitude, i.e. the difference between maximum and equilibrium range size (Fig. 3D). As we had hypothesized, a considerable Allee effect must be present for elasticity to emerge (*A* > 0 for *α* ≥ 0.02, Fig. 3D).

**Figure 2:**
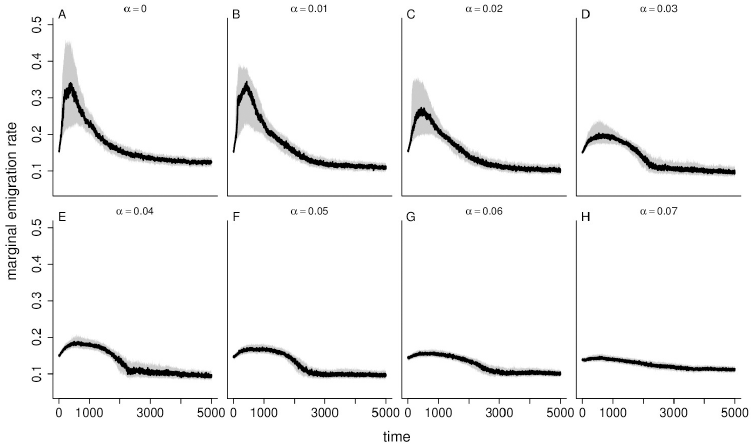
Marginal emigration rate as a function of simulation time for a gradient in dispersal mortality (*µ*). Allee effect strength increases from A to H. For parameter values see main text. The black lines show the median values of 50 replicate simulations, the shaded grey areas denote 25% and 75% quantiles.

**Figure 3:**
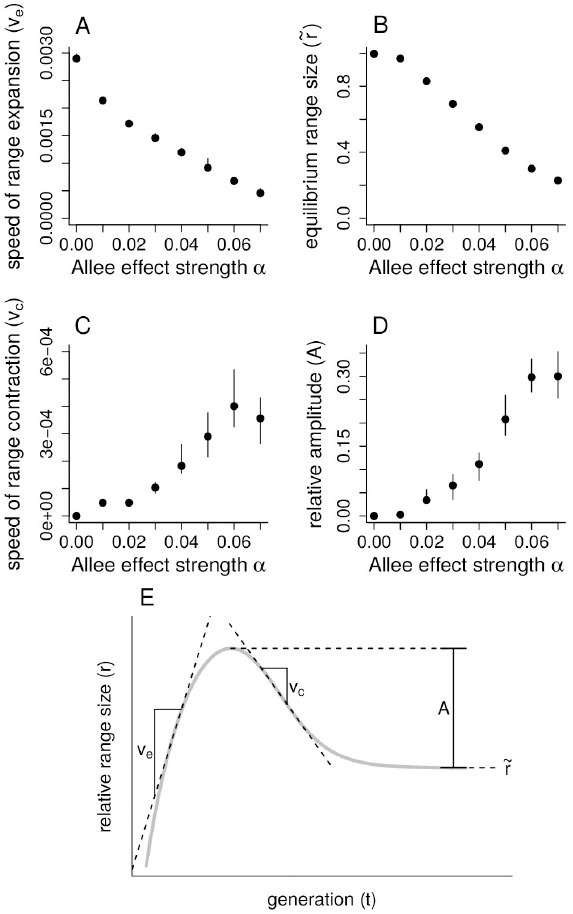
Results of the quanitative analysis of elasticity depending on Allee effect strength. Shown are A) the speed of range expansion, B) the relative range size, C) the speed of range contraction and D) the relative amplitude of the elastic range effect. Shown are median values of 50 replicate simulations, error bars denote 25% and 75%-quantiles. E) A sketch of the – for elasticity typical – relationship between relative range border position and time to illustrate the measures used in A-D. The curve was created by using eq. 3 with the following parameters: *v*_*e*_ = 0.0015, *v*_*c*_ = 0.0004, ∆_*t*_ = 2000, 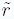 = 0.35). The meaning of the four parameters used in the analysis is denoted.

The results of simulations allowing for the evolution of dispersal distance instead of propensities show the same qualitative behavior. Elasticity can only be found in a gradient of dispersal mortality (per-step mortality) and for strong Allee effects (Supplementary material Appendix 2, Fig. A6 - A10).

The numerical analyses of mean colonizer fitness show characteristically different patterns between the various environmental gradients (Fig. 4). For the dispersal mortality gradient without Allee effects an increase in colonizer fitness deeper in the gradient can be observed (Fig. 4A), which sets this gradient apart from all the others. Increasing emigration (darker lines) results in overall lowered fitness expectations, especially for range core areas (Fig. 4A). Allee effects strongly interact with this pattern (Fig. 4B): for high dispersal rates the relationship between colonizer fitness and spatial location changes from monotonically increasing to unimodal with a rapid decrease of fitness in regions with harsher conditions (i.e. higher dispersal mortality; Fig. 4B). The lower the dispersal rate is, the sooner this decrease sets in. It is important to keep in mind that after range expansion, dispersal decreases evolutionarily at the margin in this gradient, implying a temporal change in the fitness expectations according to the results presented in Fig. 4B.

**Figure 4:**
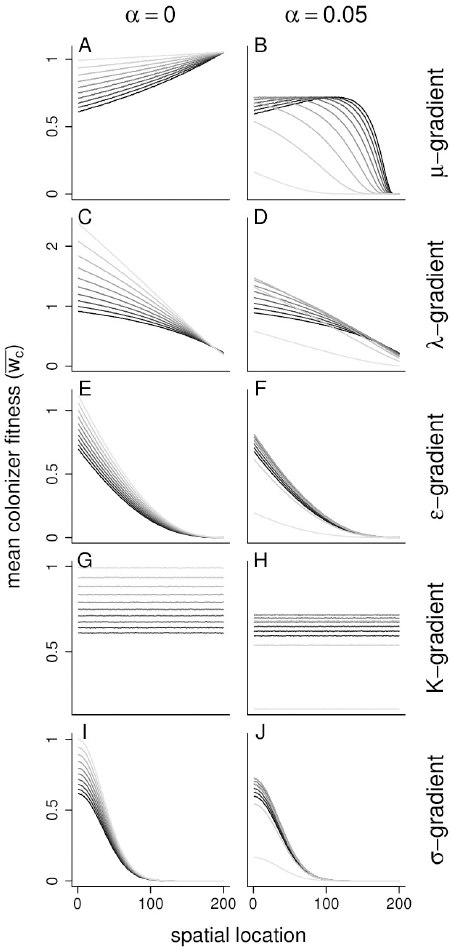
Results of the numerical analyses of mean colonizer fitness as a function of the Allee effect strength (*α*), landscape characteristics and emigration rate. Shown is the approximated geometric mean per capita offspring number to be expected for colonizers arriving at every potential *x*-location under the assumption that the species’ range ends right before each position (see main text for details). Line coloring refers to the rate of dispersal, which decreases from 0.5 (black) to 0.05 (lightest gray) in 10 equidistant steps.

This dramatic impact of Allee effects and qualitative change of the spatial distribution of colonizer fitness is strictly associated with scenarios in which (strong) range border elasticity can be observed (mortality, *µ*, and fecundity gradients, *λ*_0_, see Fig. 1, A2). For all other gradients, we either found patterns of decreasing fitness deeper in the gradient (Fig. 4C-F,I,J) or no spatial relationship in the case of the gradient in carrying capacity (Fig. 4G,H). However, for all these gradients we found the same impact of the Allee effect: whereas in its absence decreased dispersal rates (lighter colors in Fig. 4) result in consistently higher fitness, an Allee effect inverts this pattern. Importantly, no non-monotonic or interaction effects could be found in contrast to the results for dispersal mortality (Fig. 4A, B; fertility to a smaller extent, Fig. 4C, D).

## Discussion

Our results show that range border elasticity can only be observed in specific abiotic habitat gradients and in the presence of Allee effects if one focuses on intra-specific processes. Specifically, we show that an eco-evolutionary feedback loop, caused by the interplay between Allee effects, landscape structure and dispersal evolution, is necessary for ranges to show elastic behavior. As a range expansion proceeds into a fragmentation gradient (*µ*), the selective pressures at the range margin change: initially, spatial selection (more dispersive individuals being at the front in combination with fitness benefits through reduced competition; Phillips et al., 2010) leads to the evolution of increased dispersal. Yet, once the range has expanded more deeply into the gradient, dispersal is reduced by natural selection because of high habitat isolation (Fig. 2). As a result of this decrease, more and more patches turn into sinks because the reduction of the number of immigrants reduces population sizes. These small populations decrease even further due to the presence of the Allee effect. This means that from the populations’ perspective the fitness gradient gets steeper over time, ultimately resulting in a range contraction (Fig. 4B). The strength of this effect is further increased due to selection against dispersal as a result of the marginal source/sink-dynamics — a downward spiral of eco-evolutionary feedbacks leading to the observed strong range contraction.

### Eco-evolutionary feedbacks lead to range border elasticity

The fitness-relevant consequences of these non-linearly interacting effects can be seen directly in Figure 4B. High dispersal rates due to spatial selection (dark lines) lead to a fast range expansion, as colonizer fitness is initially increasing or at least not decreasing over space. At a certain point in space — defined by the interaction between gradient steepness, dispersal rate and Allee effect strength — colonizer fitness drops abruptly. Selection against dispersal due to high dispersal mortality leads to changes in the fitness profiles (depicted in light gray colors). These dynamics can only be observed in mortality gradients and to a smaller extent also in fertility gradients (Fig. 4D). All other gradients (Fig. 4E-J) do not show such an abrupt change in the spatial fitness expectations and mostly exhibit decreasing colonizer fitness early on. In fact, colonizer fitness monotonically decreases across the range for gradients in both extinction risk (Fig. 4E,F) and environmental stochasticity (Fig. 4I,J), as both factors lead to decreasing long-term population growth. The fact that colonizer fitness is not affected by the gradient in carrying capacity might seem unexpected at first. Yet, the reason is that deep in the gradient, where patch sizes are small and competition acts strongly already at small numbers of individuals, colonizer numbers are low, leveling the latter effect out. In general, the findings from our numerical analysis imply that range border elasticity cannot be found in gradients other than dispersal mortality and, to a much lower extent, growth rate. The result is in accordance with the findings by Kubisch et al. (2010), where also no elastic range behaviour could be found in gradients of patch size, growth rate and local extinction risk. While all investigated environmental gradients lead to the formation of a stable range limit, the eco-evolutionary feedback loop as described in the previous paragraph, which leads to intraspecifically caused range elasticity, is impossible in these scenarios. Especially decreasing patch size, increasing environmental stochasticity and increasing local extinction risk result in selection for increased emigration rates (Kubisch et al., 2014), which does not lead to source-sink conversion.

### Abiotic and biotic factors influencing range border elasticity

The crucial aspect of a landscape which allows for range elasticity, is not that fitness expectations of colonizers are spatially non-linear. To demonstrate this fact, we tested convex, concave and sigmoid gradient shapes with varying cline strength (Appendix 3, Fig. A11-A14). Our results are qualitatively not affected by the shape of the gradient - in the absence of Allee effects no elasticity can be observed. Hence, fitness expectations of colonizing individuals need not only to change in space, but also in time. As mentioned in the introduction this temporal change could in principle be caused by more mechanisms than the one we focus on in this study:

Abiotic changes, e.g. temporally changing climatic conditions, can also lead to elastic range dynamics. We performed additional simulations, in which we continuously worsened conditions by shifting the baselines of linear gradients. Depending on the strength of change we were able to create range elasticity in every gradient, without Allee effects (Appendix 3, Fig. A15). An example for this type of elasticity is reported in the study of Henry et al. (2013), who modeled a range shift driven by temporally shifting gradients. Similar to what we describe in this study, the populations in that scenario evolved higher dispersal during that range shift. Once climate change stopped the individuals continued to disperse further into unsuitable habitat. Subsequently, natural selection led to a reduction in dispersal at the range margin, but with a time lag between the end of climate change and the optimal adaptation of dispersal strategies. During this time lag individuals continue to disperse outside their range which leads to strong source/sink-dynamics. As Henry et al. concluded, in such a scenario the nature of the gradient is irrelevant for elasticity to occur.

Besides abiotic factors, it is also plausible that interspecific interactions can result in range elasticity. A hypothetical example would be a species expanding into a landscape, which is inhabited by a predator. If one imagines that the predator follows a Holling type III functional response this would imply that the predator species ignores the range expanding prey species as long as the latter occurs at low densities. During the course of the range expansion, however, these densities will increase, ultimately leading to a shift in the predator’s behavior, and triggering predation on the range expanding prey. This could lead to range contraction after expansion.

Both abiotic and interspecific mechanisms that can potentially result in range border elasticity are extrinsic processes — i.e. they induce an external change in the environment, which might force the focal species to contract its range. In this study, however, we focused on intrinsic, intra-specific, reasons for range elasticity. Under this premise range border elasticity can only be achieved through Allee effects in concert with decreasing dispersal towards the range margin. It is important to keep in mind that the term “Allee effect” is a phenomenological description of a variety of mechanisms ranging from sexual reproduction to social behavior (Courchamp et al., 2010).

The emergent elasticity of Kubisch et al. (2010) was also caused by an Allee effect. The authors modeled a species with sexual reproduction, thus implicitly assuming a mate-finding Allee effect. Range elasticity might even be observed in more natural gradients of fragmentation, in which not only patch isolation increases, but patch size also decreases (Fig. A5). Although lower patch sizes imply increased demographic stochasticity and thus selection for increased dispersal at the range margin, this selective force is outweighed by the strong selection for lower dispersal due to its increased costs.

### Range expansion, Allee effects and dispersal kernels

The negative relationship we find between range size and Allee effects as well as the speed of range expansion and Allee effects are in good accordance with previous theoretical studies. Using reaction-diffusion models and ordinary-differential-equation models Keitt et al. (2001) for example showed that for a wide range of biologically plausible conditions Allee effects may not only slow down invasions, but lead to the formation of stable range limits through invasion pinning (i.e. propagation failure). A good overview of the topic is provided by the literature review of Taylor and Hastings (2005), which summarizes known consequences of Allee effects for invasions and includes both theoretical and empirical work.

For reasons of simplicity and scale, we model dispersal as being restricted to surrounding populations. Yet, it is reasonable to assume that many species rather disperse varying distances, the distribution of which can be described by dispersal kernels (Clobert et al., 2012). To test the validity of our findings under the assumption of evolving dispersal distances we performed additional simulations using negative exponential dispersal kernels in analogy to Henry et al. (2013). Our results prove to be robust, as elasticity was again only found in scenarios with gradients in dispersal mortality and strong Allee effects, while being only weakly evident in fertility gradients (Supplementary Material; Figs. A6-A10).

## Conclusion

We show that elastic ranges can be caused by the eco-evolutionary interplay between dispersal evolution at the range margin, habitat fragmentation and the presence of Allee effects. If one of these conditions is not fulfilled, i.e. if the Allee effect is not strong enough or if higher dispersal is selected for at the margin than in the core (as e.g. in gradients in patch size or demographic stochasticity), range contractions after an expansion period **do** not emerge in our model. These conclusions generally hold true as long as range border elasticity is not externally triggered, for example by rapid and strong climate change or novel biotic interactions such as a change in predation. We predict that in nature, range contractions after expansions are most likely to occur in fragmentation gradients for species that reproduce sexually, show social behavior or are otherwise prone to suffer from an Allee effect.

## Acknowledgements

We thank three anonymous reviewers and Alexander Singer for thoughtful comments on an earlier version of this manuscript. EAF thanks the Eawag for funding.

